# Deciphering the enigmatic role of the amidotransferase LipL in *Bacillus subtilis*lipoic acid utilization

**DOI:** 10.1101/419234

**Authors:** Natalí B. Rasetto, Antonela Lavatelli, Natalia Martin, María Cecilia Mansilla

## Abstract

Lipoate is an essential cofactor for key enzymes of oxidative and one-carbon metabolism. It is covalently attached to E2 subunits of dehydrogenase (DH) complexes and the GcvH subunit of the glycine cleavage system. *Bacillus subtilis* possess two protein lipoylation pathways: biosynthesis and scavenging. The former requires octanoylation of GcvH, amidotransfer of the octanoate to E2s, and insertion of sulfur atoms. Lipoate scavenging is mediated by a lipoate ligase (LplJ), that catalizes a classical two-step ATP-dependent reaction. Although these pathways were thought to be redundant, a Δ*lipL* mutant, unable to transfer the octanoyl group from GcvH to the E2s during lipoate synthesis, showed growth defects in minimal media even when supplemented with this cofactor, despite the presence of a functional LplJ. In this study we demonstrated that LipL is essential to modify E2 subunits of branched chain ketoacid and pyruvate DH during lipoate scavenging. LipL must be functional and it is not forming a complex with LplJ, which suggests that these enzymes might be acting sequentially. We also show that the E2 subunit of oxoglutarate DH is a good donor for LipL amidotransfer reaction. The essential role of LipL during lipoate utilization relies on the strict substrate specificity of LplJ, determined by charge complementarity between the ligase and the lipoylable subunits. LplJ does not recognize E2 subunits without a negatively charged residue in key positions of the target protein, and thus LipL is required to transfer the lipoate to them. This model of lipoate scavenging seems widespread among Gram-positive bacteria.

## Introduction

Lipoic acid (LA) is an organosulfur compound distributed in all domains of life. Five lipoate-dependent multienzyme complexes, which are involved in oxidative and one-carbon metabolism, have been characterized (1). Pyruvate dehydrogenase (PDH) converts pyruvate into Acetyl-CoA; oxoglutarate dehydrogenase (ODH), a tricarboxylic citric acid cycle enzyme, converts oxoglutarate into succinyl-CoA; branched-chain 2-oxoacid dehydrogenase (BKDH) is an enzyme involved in branched chain fatty acids (BCFA) synthesis; acetoin dehydrogenase (ADH) acts in stationary phase of growth and converts acetoin into Acetyl-CoA. These lipoate-requiring complexes contain three catalytic subunits, E1, E2, and E3. The fifth complex, the glycine cleavage system (GCS), catalyzes the oxidative decarboxylation of glycine and is composed of four proteins, called P, H (GcvH), T and L proteins. LA is linked through an amide bond to a specific lysine residue of the lipoyl domains (LD) of the E2 and GcvH proteins, where it acts as a swinging arm transferring reaction intermediates among the multiple active sites of the enzyme complexes (2).

LA metabolism has been thoroughly characterized in the Gram-negative bacterium *Escherichia coli*. This organism has two redundant pathways for protein lipoylation: an endogenous, or *de novo* synthesis, and a scavenging pathway of the cofactor from the environment. In the first step of LA synthesis an octanoyl transferase (LipB) catalyzes the attachment of octanoate derived from fatty acid synthesis to lipoylable domains in the E2 subunits, PDH-E2 (E2p, dihydrolipoamide acetyltransferase), ODH-E2 (E2o, dihydrolipoamide transsuccinylase), and GcvH. Then, the LA synthase (LipA) catalizes the conversion of octanoyl side chain to lipoyl, by introduction of a pair of sulfur atoms (3). The scavenging pathway is directly carried out by lipoyl protein ligase A (LplA) which attaches exogenous LA to the apoproteins by a two-step ATP-dependent reaction: a) the activation of LA to lipoyl-AMP and b) the transfer of this activated lipoyl species to E2 subunits and GcvH, with the concomitant liberation of AMP (1, 4, 5).

The model Gram-positive bacterium *Bacillus subtilis* has a sole lipoate ligase, LplJ, which catalyzes the same ATP-dependent reaction as LplA, as demonstrated by *in vitro* modification of *E. coli* and *B. subtilis* apoproteins (6). However, the pathway of LA synthesis resulted more complex than that of *E. coli*. *B. subtilis* requires four protein activities to lipoylate its apoproteins *de novo*, instead of the two enzymes necessary in the Gram-negative bacterium (Fig. 1A). First, the octanoyl-acyl carrier protein (ACP):protein-N-octanoyltransferase, LipM, transfers the octanoyl moieties to GcvH (7). Then, the amidotransferase, LipL, transfers the octanoyl side chain from GcvH to the E2 subunits (8). Finally, LipA inserts sulfur atoms into C6 and C8 of the octanoyl moieties (9). It is likely that LipA can modify octanoyl-GcvH and octanoyl-E2 *in vivo*. Surprisingly, a Δ*lipL* mutant unable to transfer the octanoyl group from GcvH to the E2s during LA synthesis, also showed growth defects in minimal medium supplemented with this cofactor, albeit the presence of a functional lipoyl ligase (6). However, lipoate supplementation fully restored growth of a Δ*gcvH* mutant, which correlates with modification of all the E2 subunits (8). This result indicated that LipL, but not GcvH, is involved in the scavenging pathway. We hypothesized that LipL plays a role in lipoate scavenging by regulating LplJ activity or modulating global changes in gene expression of the target proteins, making lipoyl scavenging insufficient in its absence. In this paper, we establish the essential role of LipL in both pathways of lipoate post-translational modification, underscoring its relevance as a valid target for the design of new antimicrobial agents against Gram-positive bacteria.

**Figure 1.**
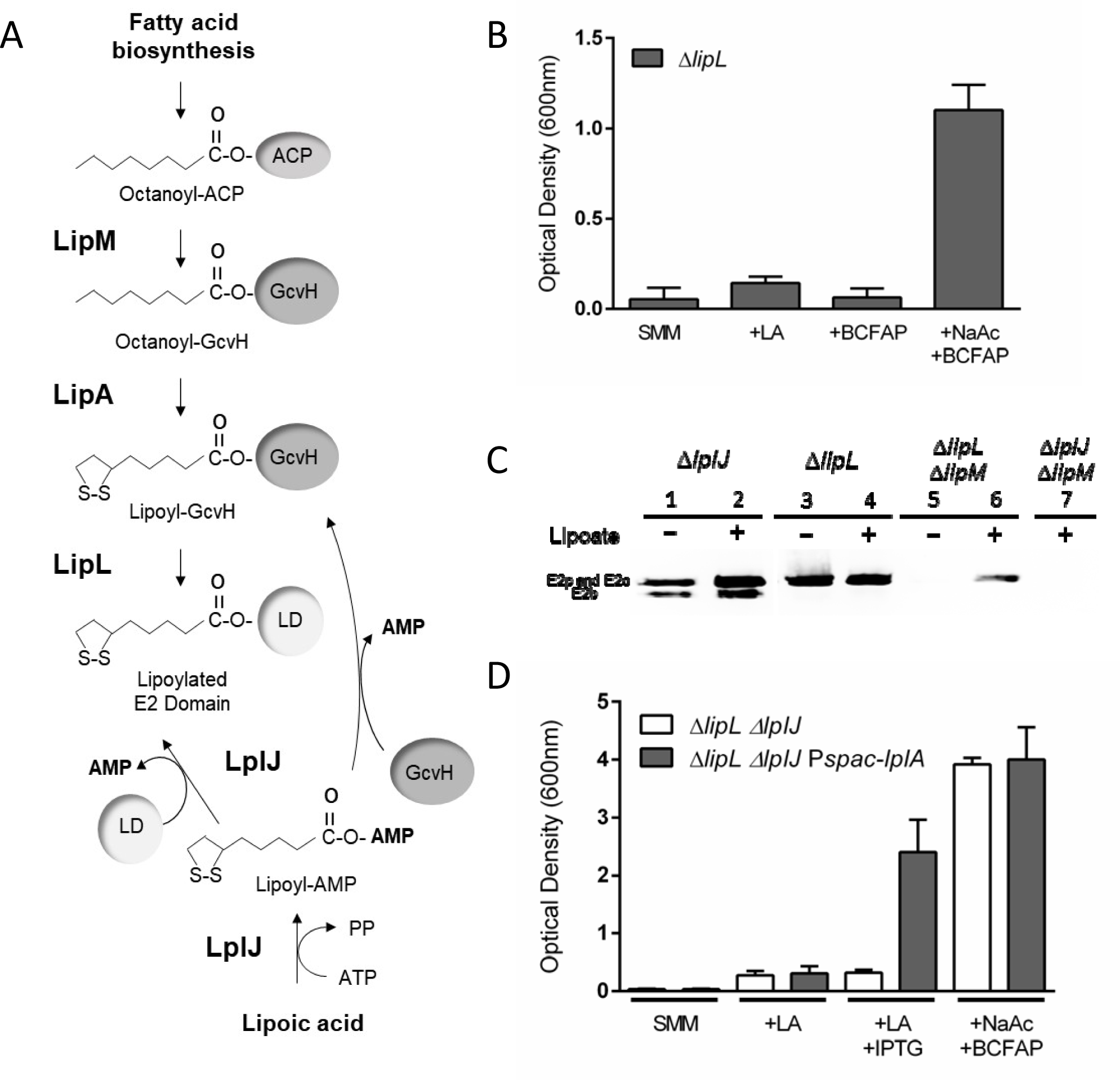
A. Current model for lipoic acid synthesis and scavenging in *B. subtilis.* During lipoic acid (LA) synthesis LipM transfers octanoic acid ligated to the acyl carrier protein (ACP), from the fatty acid biosynthesis, to the H protein of the Glycine cleavage system (GcvH). Then, LipA generates lipoyl-GcvH and LipL transfers the lipoyl group from GcvH to the lipoyl domain (LD) of the E2 subunits. Exogenous lipoate is transferred to the LDs and the GcvH subunit by LplJ in an ATP-dependent two steps reaction. **B. Growth phenotype of a *B. subtilis* mutant deficient in amidotransferase.** Strain NM51 (Δ*lipL*) was grown overnight in SMM supplemented with acetate and branched chain fatty acid precursors (BCFAP). The cultures were centrifuged and the cells resuspended in SMM or with the addition of the indicated supplements. Growth was determined by measuring the OD_600_ of the cultures at 22 h of incubation at 37°C. **C. Immunoblotting analysis of mutant strains with an anti-LA antibody.** Strains were grown overnight in SMM supplemented with acetate and BCFAP. The cells were diluted in fresh medium of the same composition with or without the addition of LA, as indicated, and grown for 22 h before analysis. **D. Effect of complementation with the *E. coli* lipoate ligase.** Strains NM67 (Δ*lipL* Δ*lplJ*) and NR001 (Δ*lipL* Δ*lplJ* P*spac-lplA*) were grown overnight in SMM supplemented with acetate and BCFAP. Cells were collected and resuspended in SMM, or with the addition of the indicated supplements. OD_600_ values of the cultures were measured after 22 h of growth at 37°C.

## Results

### *B. subtilis* lipoyl ligase LplJ requires the amidotransferase LipL to ligate lipoate to all E2 subunits

It was previously described that LplJ, the lipoyl protein ligase, is the enzyme that links lipoate to the apoproteins of *B. subtilis* (6). The lack of lipoylation of Δ*lipM* Δ*lplJ* double mutants indicates that LplJ is the sole *B. subtilis* LA salvage enzyme. Besides, expression of LplJ in *E. coli lipA lplA* cells, that are unable to synthesize and ligate LA, restored their ability to ligate LA to all *E. coli* apoproteins (6). However, modification of *B. subtilis* E2p by LplJ has not been detected *in vitro* (6). Surprisingly, *B. subtilis ΔlipL* mutants are unable to grow in SMM supplemented with LA, although a functional LplJ is present (6, Fig 1B). These observations suggest that LipL is also involved in the LA salvage process.

To determine the role of LipL in lipoate ligation to the apoproteins we performed Western blot analysis on cell extracts of *B. subtilis* mutants defective in the synthesis or scavenging pathways, grown in the presence or absence of exogenously provided lipoate. Anti-LA immunoblot of the Δ*lplJ* strain shows a wild-type pattern of lipoylated proteins: two major bands, with apparent masses of 60 and 52 kDa, were detected both, in the absence and presence of LA (Fig 1C, lanes 1 and 2). The higher molecular weight band corresponds to the E2p and E2o subunits, which run with the same apparent molecular weight in SDS-PAGE, whereas the lower molecular weight band corresponds to the E2b subunit (BKDH-E2, lipoamide acyltransferase). These results were expected since in the Δ*lplJ* strain the LA biosynthetic pathway is still functional. In contrast, immunoblot analysis of crude extracts of strain Δ*lipL* in the presence of LA showed only the higher molecular weight band, (Fig 1C, lane 4). This result denotes that LipL is required for the ligation of exogenously provided LA to, at least, the E2b subunit and explains the observed growth defect of a *lipL* mutant in SMM supplemented with LA (Fig 1B). This band is also detected when Δ*lipL* cells are grown in SMM without LA (Fig 1C, lane 3). It seems that LplJ is transferring LA from lipoyl-GcvH, which accumulates in the absence of the amidotransferase, to the E2 subunits of higher apparent molecular weight. If this is the case, a Δ*lipL* Δ*lipM* double mutant, that would not transfer the octanoyl residue to GcvH and, in consequence would not accumulate lipoyl-GcvH, will not be able to lipoylate its proteins in the absence of the exogenously provided cofactor. Our hypothesis was confirmed by the observation that there are no lipoylated proteins in extracts of Δ*lipL* Δ*lipM* cells grown in this condition (6, Fig 1C, lane 5). When LA is added to the growth medium, the higher molecular weight band can be detected (Fig 1C, lane 6), due to the ligation of the exogenous cofactor by LplJ. Besides, in extracts of Δ*lipM* Δ*lplJ* cells there aren’t any lipoylated proteins even when LA is present in the medium, due to the lack of a functional endogenous biosynthetic pathway and the absence of the lipoate ligase (Fig 1C, lane 7).

As the E2o and E2p subunits have the same apparent molecular weight in SDS-PAGE, we perform media supplementation analysis to determine if both proteins were functional in the Δ*lipL* mutant. To this end, the Δ*lipL* strain was grown in SMM supplemented with BCFAP or both sodium acetate and BCFAP, as it is already known that exogenous succinate is not a requirement for *B. subtilis* growth in SMM (9). As shown in Fig 1B, this strain is only able to grow if both sodium acetate and BCFAP are added to SMM, indicating that the PDH complex is not functional in a Δ*lipL* strain. These results also suggest that the 60 kDa lipoylated band observed in the immunoblotting analysis of this mutant (Fig 1C, lane 3 and 4), corresponds to E2o subunit.

Together, these results indicate that the lipoate ligase enzyme LplJ is essential to transfer exogenous LA to GcvH and the E2 subunits, but also requires LipL to modify E2p and E2b subunits. This differs from other lipoyl ligase enzymes, such as *E. coli* LplA, which can transfer exogenous lipoate to all E2 subunits without the requirement of an additional protein. Expression of *E. coli* LplA under the control of the IPTG-inducible promoter P*spac* in a Δ*lipL* Δ*lplJ* mutant restores growth of this strain in SMM supplemented with lipoate (Fig. 1D). Since the Δ*lipL* Δ*lplJ* double mutant is impaired in both LA biosynthesis and utilization, this result indicates that LplA can functionally bypass both pathways in *B. subtilis* without the requirement of an additional protein. As shown for LA synthesis, where a four-protein pathway is required in *B. subtilis* (6-8) instead of the two-protein lipoylation mechanism utilized by *E. coli* (10, 11), the ligation of exogenous lipoate in this Gram-positive model bacterium also follows a more complex pathway than in the Gram-negative model (4, 5).

### LipL is required for octanoic acid scavenging

In *E. coli,* the lipoyl ligase LplA is able to transfer both lipoate as well as octanoate to the apoproteins, albeit less efficiently (11). A similar behavior was observed in an *Staphylococcus aureus lipM* mutant (12). To determine whether *B. subtilis* LplJ is able to ligate exogenously provided octanoic acid and if LipL is also involved in this process, a Δ*lipM* strain was grown in SMM supplemented with octanoic acid or the combination of sodium acetate and BCFAP. Whereas octanoic acid supplementation allowed growth of the Δ*lipM* mutant to levels comparable to the wild type strain, the Δ*lipM ΔlplJ* double mutant was unable to grow in the same conditions (Fig. 2A). These results indicate that LplJ is required for the transfer of exogenous octanoate to the apoproteins and that LipM does not play a role in this process. As shown in Fig. 2B, Δ*lipL* mutant strain showed the same growth defect in SMM supplemented with octanoic acid as the observed for a Δ*lipM ΔlplJ* strain, indicating that in *B. subtilis* both LplJ and LipL are required for octanoic acid scavenging, as observed for lipoate. Since octanoate supplementation fully restored growth of a Δ*gcvH* mutant (Fig. 2B), as observed for lipoate supplementation (8), we concluded that octanoate ligation by LplJ and LipL does not require the formation of an octanoate-GcvH intermediate. This result also indicates that introduction of sulfur atoms mediated by LipA can occur either on octanoyl-GcvH or on octanoyl-E2 (at least on octanoyl-E2o, see below).

**Figure 2.**
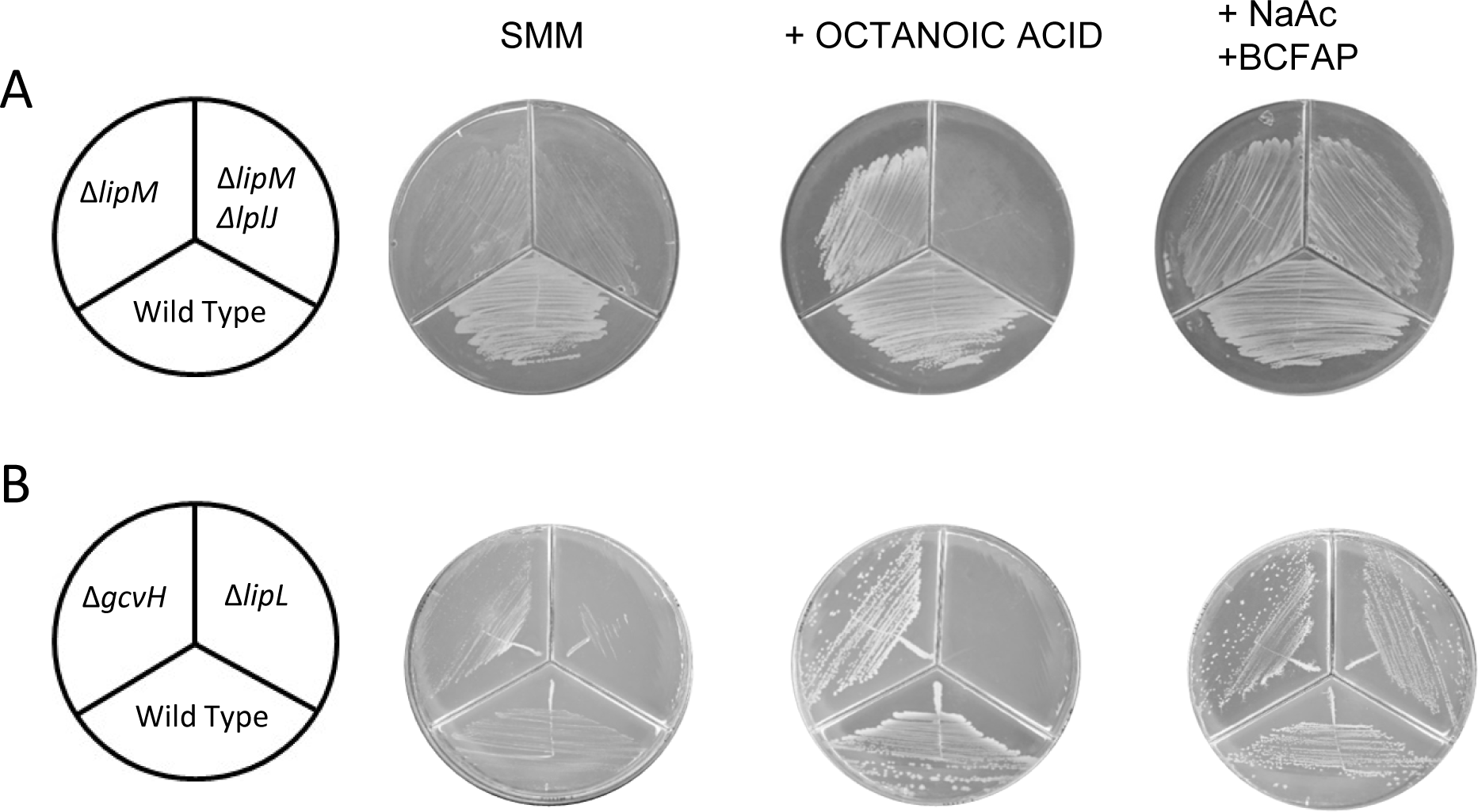
Growth of lipoic acid auxotrophs upon octanoic acid supplementation. **A.** Growth of bacterial strains JH642 (wild type), NM57 (Δ*lipM*) and NM65 (Δ*lipM ΔlplJ*). **B.** Growth of wild type (JH642), NM51 (Δ*lipL*) and NM20 (Δ*gcvH*). The strains were streaked onto SMM plates containing the supplements indicated above and incubated for 48 h at 37°C.

As reported for *E. coli* and *S. aureus*, higher concentrations of exogenous octanoic acid (125 μM) than LA (25 nM) are required to supplement growth of a Δ*lipM* mutant (Fig. S1) (10, 12). Thereby, even though *B. subtilis* LipL and LplJ are capable of transferring both exogenously provided lipoic and octanoic acid to the E2 subunits, lipoate transfer seems to be more efficient. It was recently reported that Firmicutes can uptake and incorporate extracellular fatty acids into phospholipids via the fatty acid kinase pathway (13). This pathway requires the activity of the kinase FakA and the fatty acid binding protein FakB, that act together to form acyl-phosphate. The *B. subtilis* genome contains a gene, *yloV*, with 56% similarity to *S. aureus fakA*. To determine whether octanoic acid could be directly scavenged by means of LplJ and LipL as occurs with LA, or if a FakAB-like pathway was required to synthesize phosphorylated octanoate prior to LplJ/LipL ligation to the apoproteins, we evaluated the role of *B. subtilis* YloV kinase in octanoate entry to the cell. To this end, we constructed a conditional *yloV* mutant which is also deficient in lipoate synthesis *(ΔlipM yloU∷*Pspac-*yloU*-*yloV*). This mutant was able to scavenge octanoic acid with or without IPTG-induced expression of the *yloUV* operon (Fig. S2), indicating that the FakA homologue is not required for octanoate entry into the cell, and that either free octanoate is the substrate in the scavenging pathway or another kinase is playing this role. However, as 5000-fold octanoic acid supplementation is required compared to LA, it was possible that the YloUV system was competing with LplJ for the available free fatty acid. To determine if this was the case, strain Δ*lipM yloU∷*P*spac-yloU-yloV* was grown in SMM supplemented with different concentrations of octanoic acid (0 to 500 μM). A similar growth was observed both in the presence or the absence of the inductor, reaching higher optical density with concentrations of fatty acid supplementation ≥125 μM. This result indicates that the difference in concentration requirements for exogenous octanoate and lipoate arises from a higher affinity of LplJ for lipoate. Sequence alignment of LplJ and LplA shows that the *B. subtilis* lipoate ligase contains residues predicted to form hydrophobic interactions with the dithiolane ring and the hydrophobic tail of LA, as inferred from LplA crystal structure (14). Unspecific van der Waals interactions may permit LplJ to bind LA analogues and octanoic acid, but hydrophobic interaction would be stronger when the dithiolane ring of LA is present. This might explain why LplJ has a higher affinity for lipoate than for octanoate.

### Functional LipL is required for lipoate scavenging

LplJ and LipL are both necessary for LA ligation to the E2p and E2b subunits. However, it was not clear if LipL activity was required or if LipL was fulfilling another role. To discern between these alternatives, we analyzed the growth phenotype of a Δ*lipL* mutant in which a catalytically inactive form of LipL was expressed. It was previously described that LipL residue C150 is essential for catalysis: mutagenesis of this cysteine residue resulted in loss of enzymatic activity and the inability to form an acyl-enzyme intermediate (8). Based on this evidence, an in-frame fusion of LipL C150A to the green fluorescent protein (GFP) was expressed under a xylose-inducible promoter in a Δ*lipL* mutant. As observed in Fig. 3A, the expression of the wild type version, LipL-GFP, restored the growth of the Δ*lipL* mutant even in the absence of inductor, as previously observed (6). This was probably due to the known basal expression of the P*xylA* promoter and the low levels of LipL activity required for growth. In contrast, expression of LipL C150A did not allow the growth of the Δ*lipL* mutant in the presence of LA. This result correlates with the detection of just one of the two lipoylated bands in the immunoblot corresponding to the E2p and/or E2o proteins (Fig. 3B). Although the pattern of lipoylated proteins in this strain was the same to the one of a Δ*lipL* strain, it was possible that expression of the C150A protein allowed at least modification of the E2p subunit. However, this strain was unable to grow in SMM supplemented only with BCFAP (Fig. 3A), indicating that the PDH complex is still not functional. To rule out the possibility that the observed phenotype was the result of lack of expression of the mutant version of LipL, the expression of the LipL C150A-GFP fusion protein, induced by addition of xylose, was detected by fluorescence microscopy. As shown in Fig. 3C, fluorescence was observed after the addition of the inductor to SMM, however, growth was restored only when the media was supplemented with both sodium acetate and BCFAP (Fig. 3A). These results indicate that LipL must be functional to allow E2p and E2b modification by exogenous lipoate.

**Figure 3.**
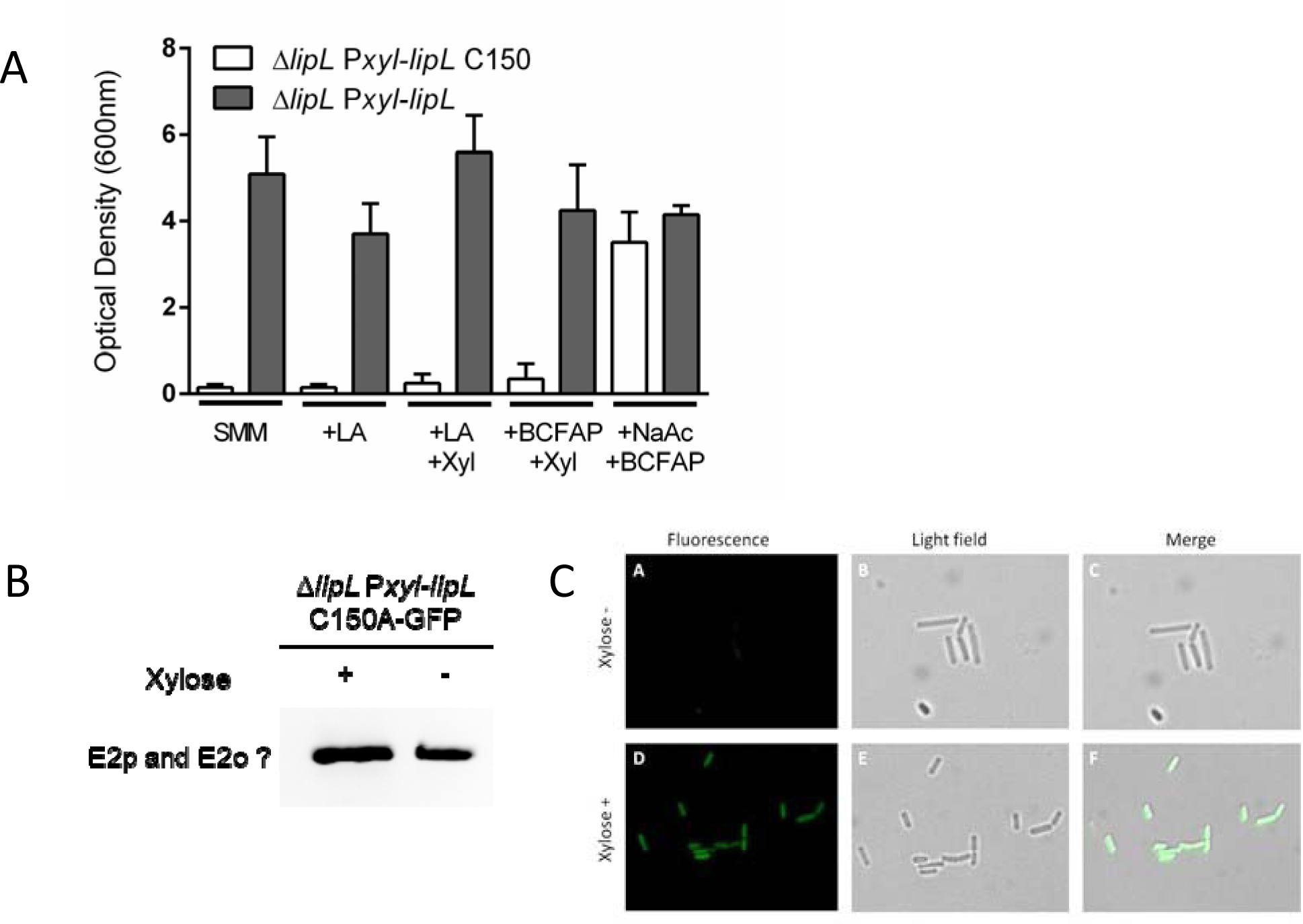
A. Growth phenotype of Δ*lipL* strain expressing LipL-C150A. Strains NR008 (Δ*lipL amyE∷lipL* C150A-GFP) and AL107 (Δ*lipL amyE∷lipL*–GFP) were grown overnight in SMM supplemented with acetate and branched chain fatty acid precursors (BCFAP). Cultures were centrifuged and cells resuspended in SMM or with the addition of supplements, in the presence or the absence of the inductor, as indicated. The OD_600_ values of the cultures were measured after 22 h of growth at 37°C. The results shown are the average of two independent experiments. **B. Lipoylated proteins of Δ*lipL* strain expressing LipL-C150A.** Strain NR008 (Δ*lipL amyE∷lipL* C150A-GFP) was grown overnight in SMM supplemented with acetate and BCFAP. Cells were diluted in fresh medium of the same composition with the addition of lipoate and with or without xylose, as indicated, and grown for 22 h before analysis. **C. Expression of LipL C150A-GFP.** The NR008 strain (NM28 *amyE∷lipL* C150A-GFP) was grown in SMM supplemented with sodium acetate and BCFAP (panels A and B). Xylose was added (0.1%) to induce the expression of LipL C150A-GFP (panels D and E). Panels C and F show the merge between fluorescence microscopy and light field microscopy in each condition.

### No interactions between LipL and LplJ are required for lipoate scavenging

We demonstrated that the E2p and E2b subunits are only lipoylated when both LplJ and a functional version of LipL are present in the cell. The dual requirement for these proteins in the utilization of exogenously provided lipoate could arise from the need of LipL and LplJ to interact forming a functional complex. Protein-protein interaction in lipoate synthesis have been proposed to occur in yeast, among Lip3, the H protein, and perhaps Lip2 and Lip5, which could be forming a lipoylation complex (15). Alternatively, LplJ and LipL could be acting sequentially. To discern between these possibilities, we used the bacterial adenylate cyclase two-hybrid system to test for LipL and LplJ interactions (16). In this system the interaction between target proteins results in the functional complementation between adenylate cyclase T18 and T25 domains. This complementation results in production of cAMP and a concomitant increase in β-galactosidase activity in *E. coli* cells. LplJ was fused to the T18 domain of the adenylate cyclase, either to the N-term and C-term, and LipL was fused to the T25 domain, also in both positions. Colonies transformed with the four possible plasmid combinations formed white colonies in LB supplemented with X-gal (Fig. 4A), even though the system successfully worked when the T18 and T25 domains were fused to interacting leucine zipper proteins (Fig. 4B). These results suggest that LipL and LplJ do not interact *in vivo* and thus, these proteins might be acting sequentially during LA scavenging.

**Figure 4.**
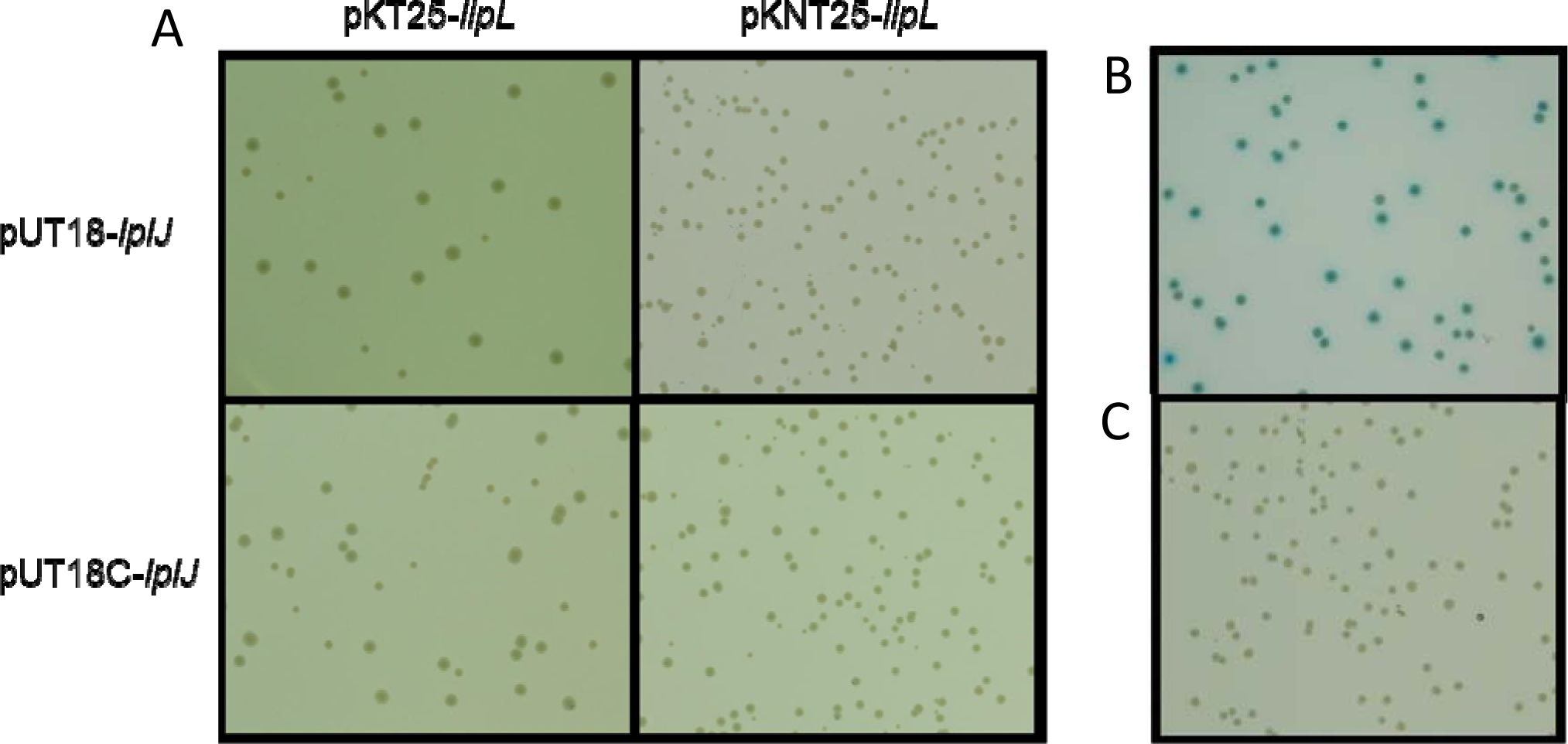
Interaction of LipL and LplJ in a bacterial adenylate cyclase two-hybrid assay. BTH101 host cells co-transformed with a pair of plasmids were selected in LB medium with X-gal and incubated at 30°C for 24 h. **A.** Plasmids containing adenylate cyclase domain T18 fused to LplJ and T25-fused to LipL **B.** T18 and T25 domains fused to leucine zipper (pUT18C-zip and pKNT25-zip), used as positive control. **C.** Empty plasmids (pKT25 and pUT28), used as negative control.

### The role of LipL in lipoate utilization

We have demonstrated that the *B. subtilis* lipoate ligase requires the presence of a functional amidotransferase to ligate lipoate to E2p and E2b. GcvH, the only known substrate of LipL and essential during *de novo* biosynthesis, resulted dispensable for lipoate utilization (8). Therefore, we hypothesized that another lipoylated protein was acting as source for the amidotransfer reaction during lipoate utilization. In Western blot assays of extracts of *lipL* mutants grown in the presence of lipoate we observed the band of lipoylated E2o (Fig.1 and 3). This protein has not been previously described as a lipoate donor in the biosynthesis pathway, because LipM only transfers octanoate from octanoyl-ACP to GcvH. However, we reasoned that lipoyl-E2o might be a good source of cofactor for LipL amidotransfer reaction during the lipoate scavenging pathway, in the absence of lipoyl-GcvH. Since it was demonstrated that *L. monocytogenes* LipL catalizes a reversible reaction (17), it is possible that the lipoylation relay uses E2o as a LA donor to transfer the cofactor to E2b and E2p. To test this hypothesis, we constructed a Δ*gcvH* Δ*odhB* strain (being *odhB* the gene encoding E2o). This double mutant was unable to grow in SMM even when supplemented with LA (Fig. 5A), and its E2s are not lipoylated in these growth conditions (Fig. 5B). These results indicate that the Δ*gcvH* Δ*odhB* double mutant lost the ability to utilize exogenous lipoate, even when wild type LplJ, LipL, and the essential lipoyl-dependent E2p and E2b are present in the cell. As expected, an Δ*odhB* strain is able to synthesize LA, and thus grows and lipoylates its apoproteins in SMM (Fig. 5). All these results allowed us to propose a model of LA biosynthesis and utilization in *B. subtilis*, where the amidotransferase LipL plays a central role in both pathways (Fig. 6).

**Figure 5.**
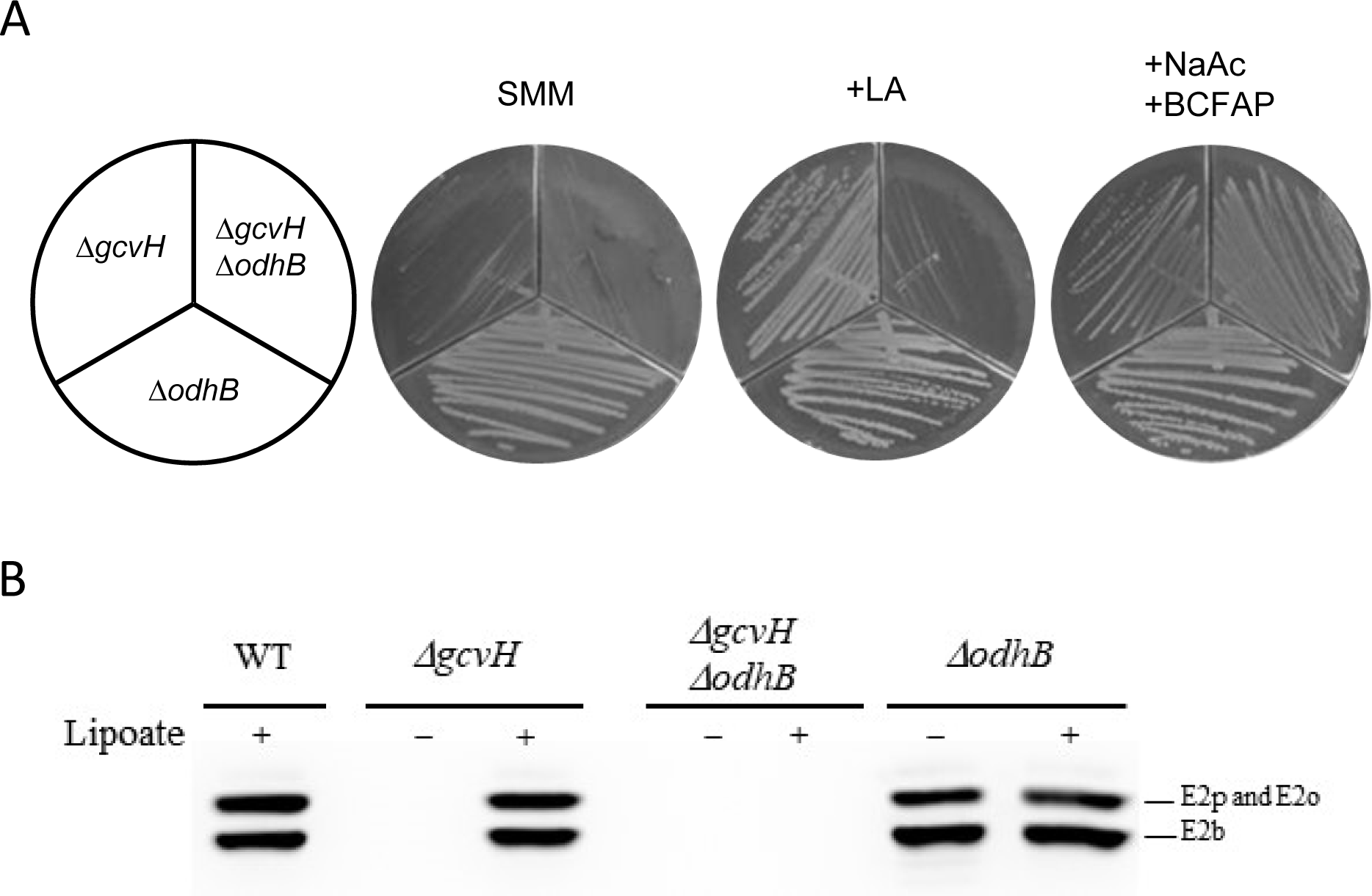
Role of oxoglutarate dehydrogenase in lipoic acid scavenging. **A.** Growth of bacterial strains CM57 (Δ*odhB*), NM20 (Δ*gcvH*) and CM56 (Δ*gcvH* Δ*odhB*). Strains were streaked onto SMM plates containing the supplements indicated above and incubated for 48 h at 37°C. **B.** Immunoblotting analysis of mutant strains with an anti-lipoic acid antibody. The strains were grown overnight in SMM supplemented with acetate and BCFAP. Cells were diluted in fresh medium of the same composition with or without the addition of lipoic acid (LA), as indicated, and grown for 22 h before analysis.

**Figure 6.**
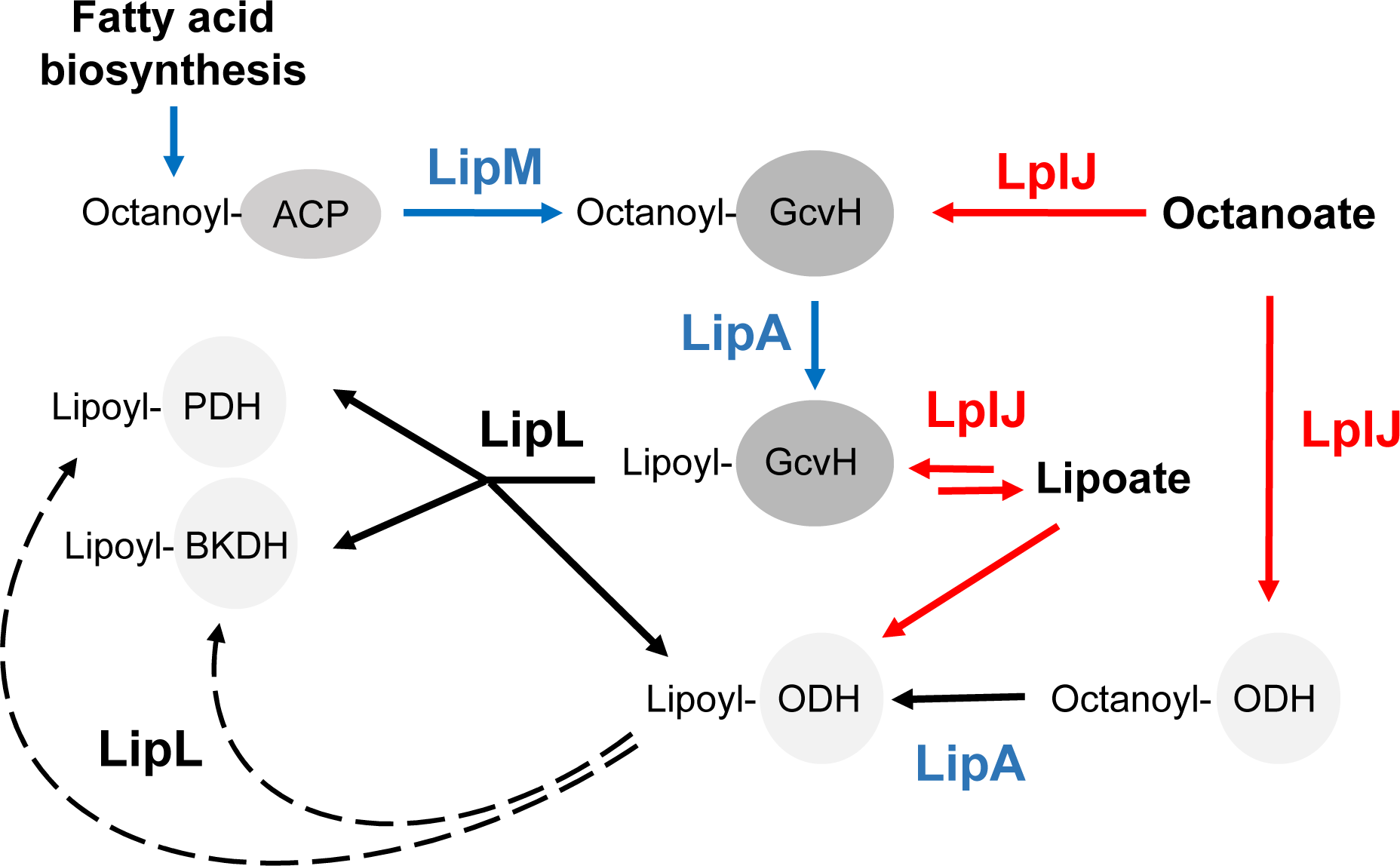
The role of LipL in lipoate and octanoate utilization. . Blue arrows: lipoic acid biosynthesis. Red arrows: lipoic and octanoic acid salvage. Black arrows: common steps. In the absence of lipoate biosynthesis the amidotransferase can transfer the lipoyl moiety from E2o to the others E2 subunits (dashed arrows). If LipL is absent, E2p and E2b cannot be modified either by the endogenous or the exogenous lipoylation pathways.

### LplJ requires a specific glutamate residue in the target apoprotein

We demonstrated that LplJ can only ligate lipoate to GcvH and to E2o, while E2p and E2b are not modified by this enzyme. We hypothesized that LplJ substrate specificity could be due to the orientation of the lipoylable lysines in GcvH and E2o, that allows a convenient interaction between the ligase and these subunits. The orientation of the corresponding lysines on the other E2 apoproteins would not favor LplJ interaction. To corroborate our hypothesis, we aligned E2o-LD I-Tasser generated models with E2p-LD, E2b-LD and GcvH models. As shown in Fig. 7, E2o and E2p lysine residues have different orientations, which would account for their differences in lipoylation. However, even though the E2b conserved lysine has the same orientation than the corresponding E2o residue, LplJ is not able to ligate lipoate to E2b. Besides, conserved lysines from E2o and GcvH, both lipoylated by LplJ, have different orientations. We conclude that orientation of the lipoylable lysine residue is not a determinant during lipoate ligation by LplJ.

**Figure 7.**
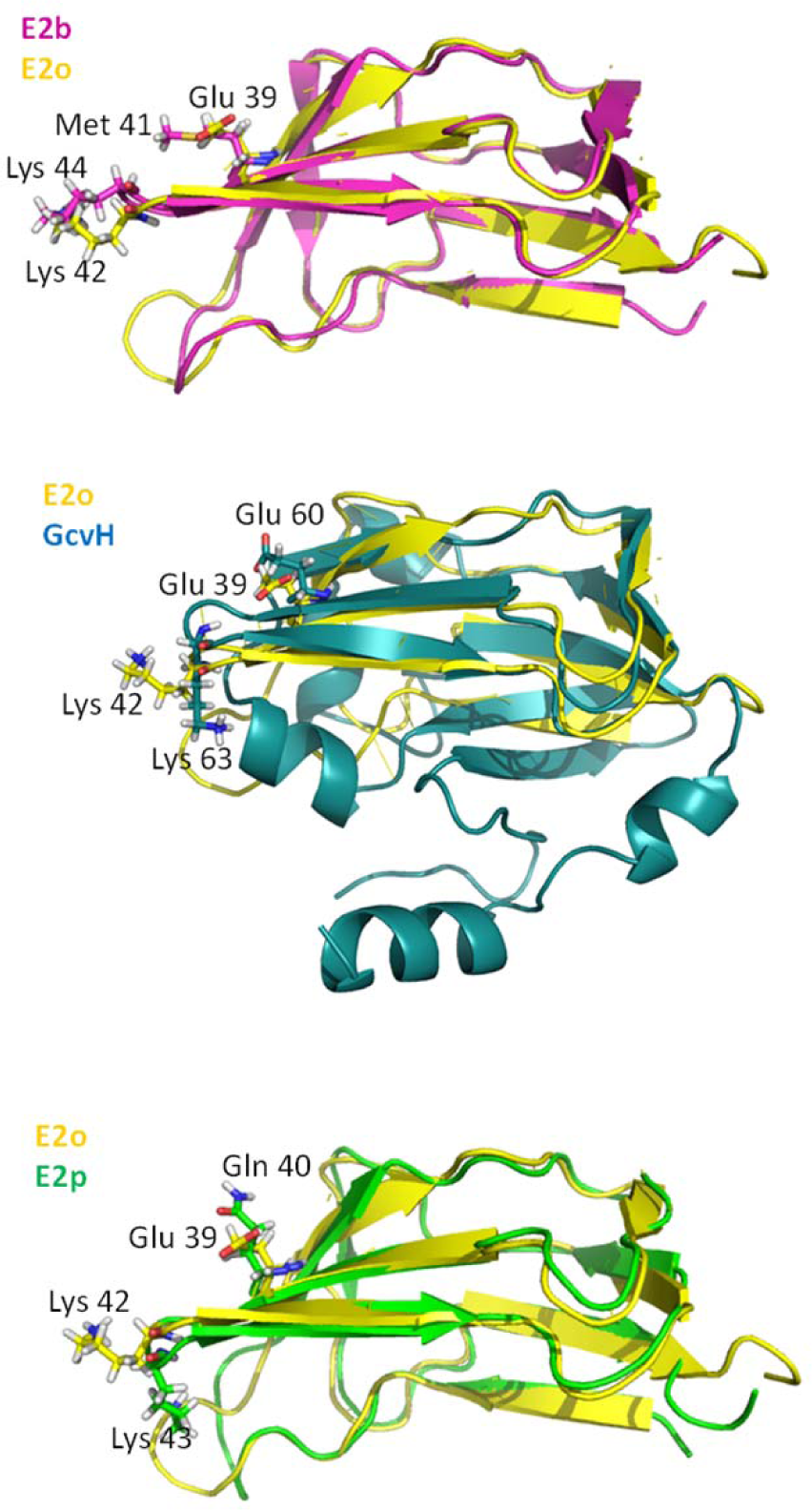
Superposition of LD-E2 and GcvH models. Ribbon representation of LD-E2b (pink), LD-E2p (green), and GcvH (blue) aligned with LD-E2o (yellow). Lipoylable lysine residues and the amino acids located at three residues on the N-terminal side from this residue (glutamate, methionine and glutamine), are shown as stick representation and labeled accordingly. The figure was generated using I-Tasser (41) and Pymol (42).

To try to identify amino acid residues that would be involved in LplJ substrate specificity, we aligned E2 LD and GcvH sequences from *B. subtilis* and *E. coli*. As shown in Fig. 8, the E2 subunits that can act as substrates for LplJ have a conserved glutamate residue, located three residues to the N-terminal side from the lipoylable Lys residue. However, E2p and E2b have a glutamine or a methionine residue instead. Modeling of the complexes between LplA of *Thermoplasma acidophilum* and E2p from *Azotobacter vinelandii* or GcvH from *Thermus thermophilus* have been performed. These models predicted interactions between conserved acidic residues from the receiver apoproteins, located close to the lipoylable Lys, and basic residues from LplA, through hydrogen bond unions (18). The apoproteins that can be modified by LplJ contain a negatively charged residue in this conserved position (Fig. 8, Glu in red), while E2p and E2b contain uncharged residues. Thus, the substrate specificity of the ligase for the lipoate acceptor protein could be determined by charge complementarity between the ligase and the lipoylable subunits. To determine if the negative charge of E2o Glu^39^ is essential during lipoylation by LplJ, we generated a mutant subunit where this glutamate residue was replaced by Gln (E2o-E39Q). As previously shown, the double mutant strain Δ*gcvH* Δ*odhB* is unable to grow in SMM supplemented with LA (Fig. 5), due to the lack of an appropriate protein recipient for LplJ ligation. When this strain was transformed with a plasmid that allows expression of the E2o wild type copy, it recovered its ability to attach LA and hence to grow in SMM in the presence of the cofactor (Fig. 9). However, when the mutant copy E2o-E39Q was expressed, the bacterial strain was unable to ligate exogenous LA, demonstrating that Glu^39^ is indeed essential for the ligation reaction carried out by LplJ (Fig. 9).

**Figure 8.**
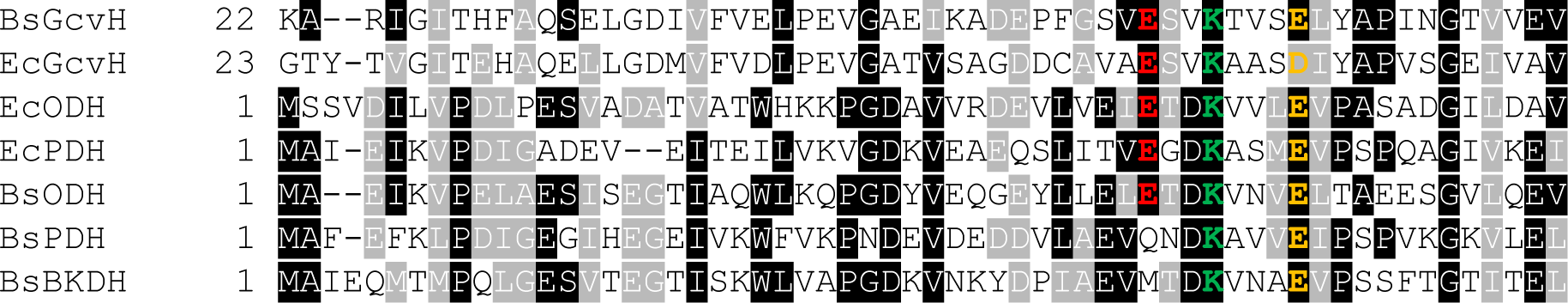
Sequence alignment of *B. subtilis* and *E. coli* E2-LD and GcvH subunits. Identical residues are shown highlighted in black and similar residues are highlighted in grey. The conserved lipoylable lysine residue is shown in green. The glutamate residue, predicted to interact with LplJ positive charges, is shown in red. The other conserved negatively charged residue that would stabilize the interaction with LplJ is shown in yellow. Bs: *B. subtilis*; Ec: *E. coli.*

**Figure 9.**
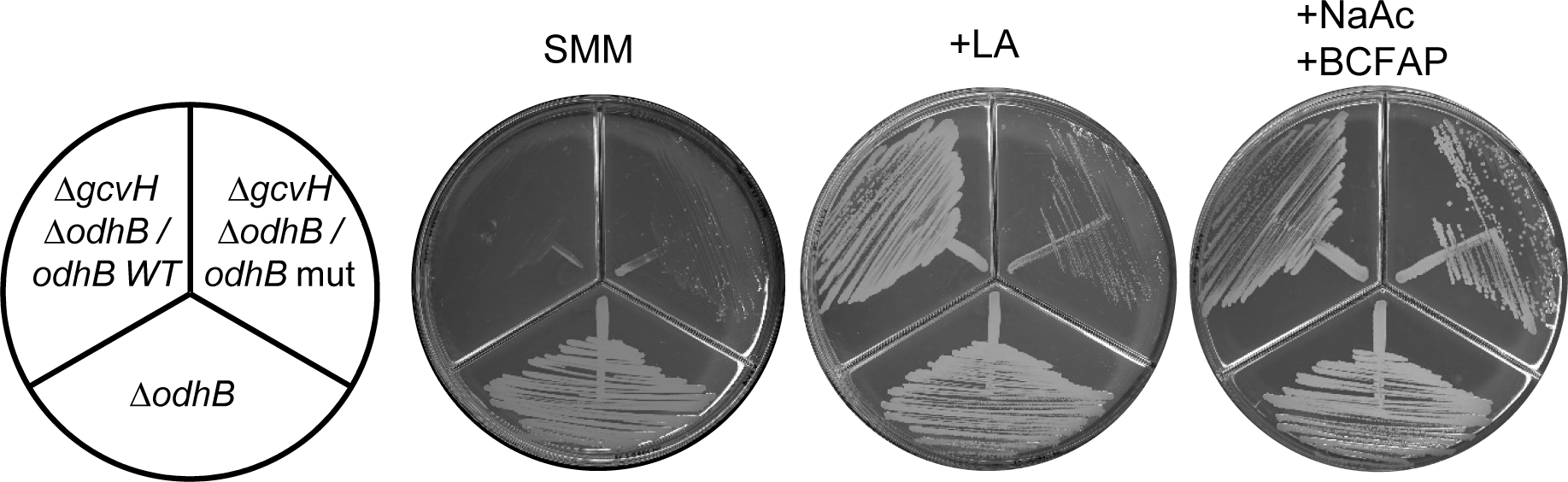
Relevance of E2o Glu^39^ for LplJ reaction. Growth of bacterial strain CM56 (Δ*gcvH* Δ*odhB*) transformed with plasmids that allow expression of either wild type E2o or E2o-E39Q copy. Strain CM57 (Δ*odhB*) was used as a positive control for growth. The strains were streaked onto SMM plates containing xylose and the supplements indicated above, and incubated for 48 h at 37°C.

## Discussion

Protein lipoylation is a post-translational modification present from bacteria to mammals. It is essential for the activity of 2-oxoacid dehydrogenase complexes and the GCS. Different organisms have evolved diverse strategies for protein lipoylation: some of them synthesize the cofactor, others utilize LA acquired from the environment, and others encode both pathways. To add to the complexity of protein lipoylation pathways, modification of apoproteins using exogenous lipoate occurs by several mechanisms. In *E. coli* LA is transferred to apoproteins in an ATP-dependent process by the lipoyl protein ligase A (LplA). In Gram-positive bacteria, the scavenging pathways are even more diverse. The model bacterium *B. subtilis* has a sole lipoate ligase, LplJ, which catalyzes the same ATP-dependent reaction as LplA (6). In contrast, the Gram-positive pathogenic bacterium *Listeria monocytogenes* possess two lipoyl ligases. While LplA1 is required for intracellular growth and can use host-derived lipoyl-peptides as substrates, LplA2 utilizes only free lipoate and is dispensable for intracellular growth (19). Additionally, LplA1 has a tight substrate specificity as it only ligates lipoate to GcvH (17). Modification of E2 LDs requires the activity of the amidotransferase, LipL, which utilizes lipoylated GcvH as substrate (17). *Staphylococcus aureus* also has two ligases: LplA1 and LplA2. LplA1 is the primary LA salvage enzyme in broth culture, while either LplA1 or LplA2 stimulate bacterial survival within macrophages in a manner dependent on exogenous LA provision (12). *In vitro* studies determined that these ligases target different LD-containing proteins: LplA1 is able to modify GcvH and E2o, while LplA2 modifies only oxoacid dehydrogenase E2 subunits (20). As expression of LplA2 is limited in broth culture, modification of E2b and E2p in this condition requires the transfer of the lipoyl moiety from lipoyl-GcvH to the apoproteins, mediated by LipL (12).

*B. subtilis* relays on two pathways for protein lipoylation, but they are not completely redundant: growth and lipoylation phenotypes observed in a Δ*lipL* mutant pointed out that the amidotransferase is implicated in both the scavenging and the *de novo* biosynthetic pathway of the cofactor. Although LplJ is able to modify all the *E. coli* E2s, we observed that in *B. subtilis* it can only transfer exogenous lipoate to GcvH and E2o. LplJ requires the presence of LipL to modify E2p and E2b subunits, which correlates with *in vitro* evidence of LplJ lipoylating GcvH but not E2p (8). Until this study the exact role of this amidotransferase during LA scavenging remained elusive.

During the *de novo* biosynthetic pathway the requirement for LipL is due to the specificity of LipM for GcvH. Based on this data, it could be inferred that LplJ modifies GcvH and then LipL catalyzes the amidotransfer reaction from GcvH to E2 subunits, as already described in *S. aureus* and *L. monocytogenes* (12, 17). However, it was reported that a *B. subtilis ΔgcvH* mutant is able to grow in SMM with lipoate as supplement, showing a strong lipoylation of the E2 subunits (6). Therefore, we concluded that GcvH is not an essential intermediate during LA scavenging. An alternative hypothesis was that LipL and LplJ formed a complex, as it was proposed to occur with the proteins Lip3, the H protein and probably Lip2 and Lip5, involved in LA synthesis in yeast (15). However, we demonstrated via two-hybrid assay that LipL and LplJ are not interacting, indicating that these enzymes are probably functioning in successive enzymatic steps. Evidence in support to this result stems from the finding that LipL must be functional during LA scavenging process.

Considering that GcvH is not required during lipoate scavenging, that LplJ can only modify E2o subunits and GcvH, and that LipL activity is necessary to lipoylate the E2p and E2b subunits, we propose that the scavenging pathway could consist of successive steps that include the dihydrolipoamide transsuccinylase (E2o). Initially, LplJ would activate LA and modify the E2o and GcvH subunits. In a subsequent reaction, LipL would catalyze the amidotransfer reaction from lipoyl-E2o and/or lipoyl-GcvH to E2b and E2p subunits. The growth and lipoylation phenotypes from a Δ*gcvH ΔodhB* strain, indicated that this double mutant is unable to utilize exogenous lipoate, even when wild type LplJ, LipL, and the lipoyl-dependent E2p and E2b are present in the cell. The need for LipL activity during LA scavenging would be due to the inability of the lipoyl ligase to utilize E2p and E2b as substrates. A comparison of the amino acids sequences surrounding the lipoylation site of GcvH and E2-LD from *B. subtilis* and *E. coli* highlighted key differences. While the proteins that can be modified by LplJ possess a Glu residue located 3 positions to the N-terminal side of the lipoylable Lys, a non-polar or uncharged residue was found in *B. subtilis* E2p and E2b. Replacement of this acidic residue by a nonpolar one in E2o (E2o-E39Q) precluded its lipoylation by LplJ, and resulted in the inability to utilize exogenous lipoate in a Δ*gcvH* background (Fig. 9). Thus, when an acidic residue is present in this position of the apoproteins, LplJ is able to lipoylate their substrates. However, the absence of this negative charge interferes with LplJ recognition. Based on structural analysis, similar interactions had been proposed between Glu residues situated in equivalent positions of E2o and E2p from *E. coli* with Gly^74^ of LplA, through a hydrogen bond (21), and from the modeled complex between *T. acidophylum* LplA and E2p from *A. vinelandii*, or GcvH from *T. thermophilus* (18). It was suggested that these residues would participate in the recognition of the apoproteins by the ligase. In this study we have demonstrated their essentiality for the reaction to proceed *in vivo*.

As *B. subtilis* E2p and E2b have a glutamine and a methionine instead of glutamate in the position equivalent to E2o-Glu^39^ they require the LipL amidotransfer activity to be modified by exogenously provided lipoate. Interestingly *B. subtilis* and mammalian amidotransferases show significant differences in substrate recognition. These differences could be a consequence of the lack of significant sequence similarity between both proteins or to differences in their mechanisms of reaction. It had been demonstrated that a Glu residue located in equivalent positions of bovine liver mitochondria E2 subunits is essential for the lipoate attachment reaction using bovine LIPT1 (22), while the presence of this charged residue does not seem to be required for LipL. However, the assayed reaction of LIPT1 corresponds to the formerly believed lipoyltransferase activity of this protein: transference of the lipoyl moiety from lipoyl-AMP to apo-LDs. Amidotransferase activity of purified human LIPT1, using E2p as receptor apoprotein, has been recently demonstrated (23). It remains to be determined if human E2b, that contains a Gln residue instead of Glu located 3 residues to the N-terminal side of the lipoylation site, could also act as a substrate in this reaction. Alternatively, another enzyme could be required for lipoamide acyltransferase modification. This is likely to be the case as LIPT1 deficiency in humans greatly alters E2p and E2o lipoylation, but E2b modification is only partly affected (24).

A similar lack of recognition of E2 subunits by the lipoyl ligases was described in *S. aureus* and *L. monocytogenes* (12, 17). We found that E2o from *S. aureus* and GcvH from both bacteria, which can be modified by LplA1 ligases, contain the conserved Glu residue located 3 positions to the N-terminal side of the lipoylable Lys. As expected, the E2 apoproteins that require LipL activity to get lipoylated contain uncharged or non-polar residues occupying these positions (Fig. S3). It is interesting to note that LplA1 from both bacteria have higher sequence similarity to *B. subtilis* LplJ than LplA2 (*S. aureus* LplA1 and LplA2 57% and 39% identity; *L. monocytogenes* LplA1 and LplA2 65% and 51% identity, respectively) and they share the same recognition requirements (17, 20). The growth phenotype and lipoylation pattern of *S. aureus* Δ*lipL* mutants (12) indicate that the amidotransferase would be performing the same role in lipoate scavenging as its *B. subtilis* orthologue.

It was previously postulated that GcvH provides an environment that facilitates the LipL reaction and that the E2-LD lack this property (25). However, in this work we demonstrate that E2o is a good substrate for LipL. Therefore, LipM ability to transfer octanoate only to GcvH, and not to any E2, would be the cause of the lipoate relay during LA synthesis in *B. subtilis*. This study demonstrated that LipL is more flexible in substrate recognition than LipM and LplJ. Further work would be required to define the determinants of this substrate specificity. LipM is able to modify all *E. coli* E2 subunits (7), but is unable to modify any *B. subtilis* E2, *Homo sapiens* E2p and GcvH2, GcvH3 and GcvH5 of *Aquifex aeolicus*. However, most of them contain the Glu residue essential for the recognition of the ligases (6, 25).

Based on our results, we propose a model for lipoate biosynthesis and utilization in *B. subtilis*, where LipL plays an essential role in both pathways transferring LA to the essential E2p and E2b, using either GcvH or E2o as donors (Fig. 6). The similarities between protein lipoylation requirements in *B. subtilis, L. monocytogenes* and *S. aureus* suggest that this dual role of LipL is conserved among Gram-positive bacteria. Due to the involvement of LA metabolic proteins in pathogenesis, multidrug resistance and intracellular growth of pathogens (26–29) the finding of essential proteins implicated in LA metabolism would provide new targets for antimicrobials. Besides, as LipL has no significant primary sequence homology with human proteins, we propose that this enzyme would be a good target for the design of new antimicrobial agents.

## Experimental procedures

### Bacterial strains and growth conditions

Bacterial strains used in this work are listed in Table I. *B. subtilis* strains are derivatives of JH642. *E. coli* and *B. subtilis* strains were routinely grown in Luria Bertani (LB) broth (30). Spizizen salts (31), supplemented with 0.5% glucose, trace elements and 0.01% each of tryptophan and phenylalanine were used as the minimal medium (SMM) for *B. subtilis*. SMM was supplemented with 50 nM or 0.5 mM DL-α-LA, 10 mM sodium acetate and 0.1 mM each BCFA precursor (BCFAP, isobutyric acid, isovaleric acid and 2-methylbutyric acid), as indicated. Xylose was added to 0.1% and isopropyl β-D-thiogalactopyranoside (IPTG) was added to 1 mM as required. Glycerol (0.5%) was used as a carbon source instead of glucose for the experiments involving gene expression under the control of the xylose-inducible promoter (P*xylA*). Antibiotics were added at the following concentrations: sodium ampicillin (Amp), 100 μg/ml; chloramphenicol (Cm), 5 μg/ml; kanamycin sulfate (Km), 5 μg/ml for *B. subtilis* or 50 μg/ml for *E. coli*; streptomycin (Str), 100 μg/ml; erytromycin (Em), 0.5 μg/ml; lincomycin (Lm), 12.5 μg/ml and spectinomycin sulfate (Sp), 50 μg/ml.

**Table 1.** Bacterial strains

| Strain | Relevant characteristics<sup>a</sup> | Source or reference |
| --- | --- | --- |
| ***B. subtilis*** |  |  |
| AL106 | *lipM::Km<sup>r</sup> yloU::Pspac-yloU-yloV* | This study |
| AL107 | *lipL::Km<sup>r</sup> amyE::lipL-GFP* | This study |
| CM28 | *lipM::Km<sup>r</sup> lipL::Sp<sup>r</sup>* | (6) |
| CM56 | *gcvH::Km<sup>r</sup> odhB::Sp<sup>r</sup>* | This study |
| CM57 | *odhB::Sp<sup>r</sup>* | This study |
| JH642 | *trpC2 pheA1* | Laboratory stock |
| NM107 | *lplJ::Sp<sup>r</sup> sacA::Pspac-lplA* | This study |
| NM20 | *gcvH::Km<sup>r</sup>* | (8) |
| NM28 | *lipL::Km<sup>r</sup>* | This study |
| NM51 | *lipL::Sp<sup>r</sup>* | (6) |
| NM57 | *lipM::Km<sup>r</sup>* | (6) |
| NM60 | *lplJ::Sp<sup>r</sup>* | (6) |
| NM65 | *lplJ::Sp<sup>r</sup> lipM::Km<sup>r</sup>* | (6) |
| NM67 | *lplJ::Sp<sup>r</sup> lipL::Km<sup>r</sup>* | (6) |
| NR001 | *lplJ::Sp<sup>r</sup> lipL::Km<sup>r</sup> sacA::Pspac-lplA* | This study |
| NR008 | *lipL::Km<sup>r</sup> amyE::lipL C150A-GFP* | This study |
| ***E. coli*** |  |  |
| BTH101 | F<sup>-</sup>, *cya-99*, *araD139*, *galE15*, *galK16*, *rpsL1 (Str<sup>r</sup>)*, *hsdR2*, *mcrA1*, *mcrB1*. | (16) |
| DH5α | *supE44* *thi-1*<br>*ΔlacU169(Φ80lacZΔM15)endA1*<br>*recA1 hsdR17 gyrA96 relA1 trp6*<br>*cysT329::lacinm<sup>λ</sup> p1(209)* | Laboratory stock |
| W3110 | F<sup>-</sup> lambda<sup>-</sup> IN(rrnD-rrnE)1 rph-1 | (43) |

### Genetic techniques

*E. coli* competent cells were transformed with supercoiled plasmid DNA using the calcium chloride procedure (32). Transformation of *B. subtilis* was carried out by the method of Dubnau and Davidoff-Abelson (33). The *amy* phenotype was assayed with colonies grown for 48 h in LB starch plates by flooding the plates with 1% I2-KI solution (34). Under these conditions, *amy*^*+*^ colonies produced a clear halo, whereas *amy*^*-*^ colonies gave no halo.

### Plasmids and strains construction

In all cases DNA fragments were obtained by PCR using the oligonucleotides described in Table II. Chromosomal DNA from *B. subtilis* JH642 was used as a template. Sanger sequencing was used to corroborate the identity and correct sequence of all the cloned fragments.

**Table 2.**
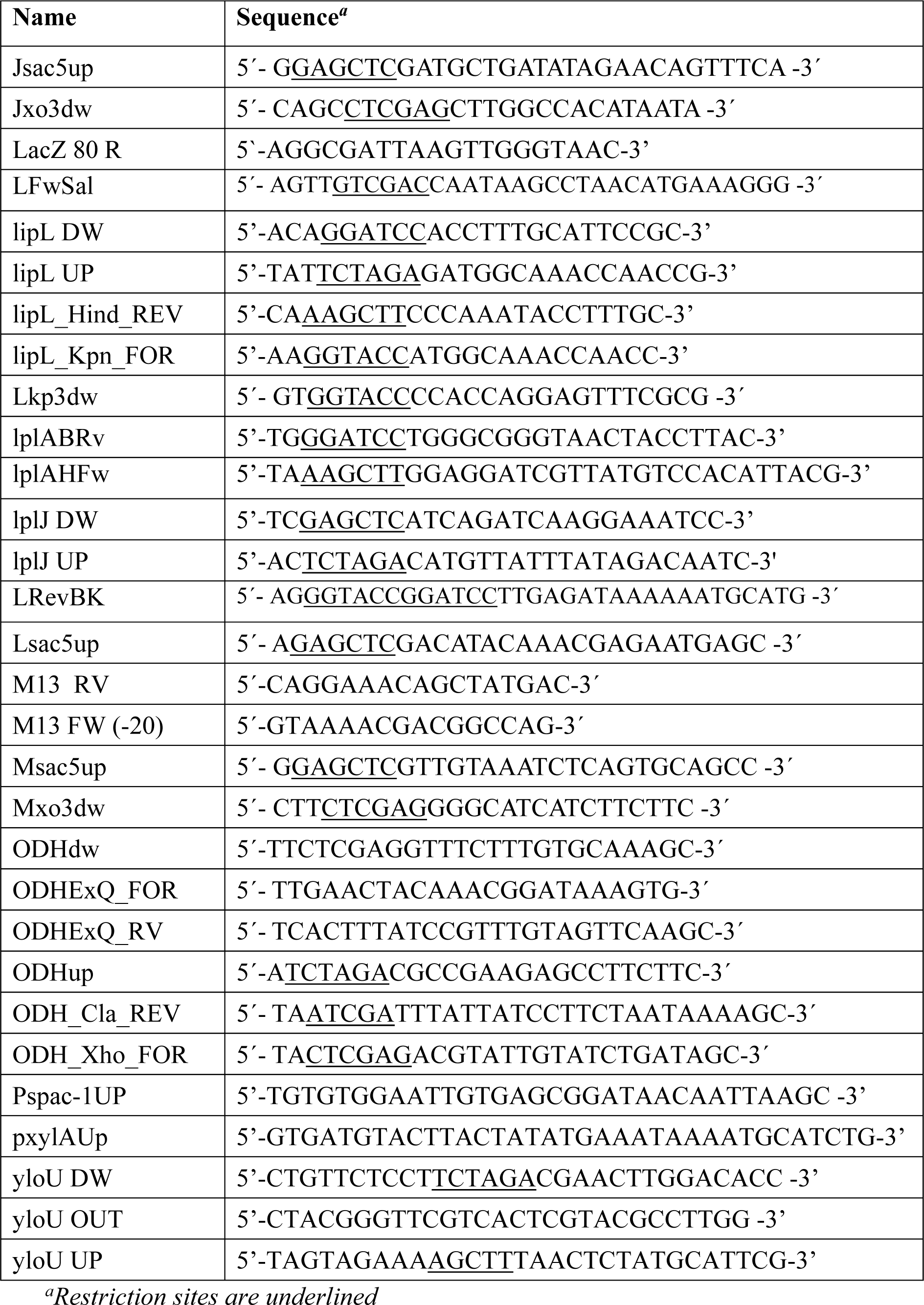
Oligonucleotide primers

**Table 3.** Plasmids used in this study

| Plasmid | Relevant characteristics<sup>a</sup> | Source or reference |
| --- | --- | --- |
| pAL26 | pSG1154 containing *lipL* gene between *KpnI*-*HindIII* | This study |
| pAL34 | pSG1154 containing *odhBE39Q* alele between *XhoI* and *ClaI* sites | This study |
| pAL35 | pSG1154 containing *odhB* gene between *XhoI* and *ClaI* sites | This study |
| pAL36 | pHPKS containing *odhBExQ* gene under a xylose inducible promoter | This study |
| pAL41 | pHPKS containing *odhB* gene under a xylose inducible promoter | This study |
| pBluescript KS | *E. coli* cloning vector, Amp<sup>r</sup> | Stratagene |
| pCM1103 | pJET1.2/blunt containing *odhB* | This study |
| pCM1104 | pJET1.2/blunt containing *odhB* interrupted with a Sp<sup>r</sup> cassette | This study |
| pDH88 | Integrative vector with polylinker under Pspac, Sp<sup>r</sup> | (37) |
| pFM24 | Integrative vector derived from pDH88 cloned with a fragment containing the *yloU* 5' region | F. Machinandiarena (unpublished) |
| pGES40-LipL | pGES40 containing *lipL* with its ribosome binding site cloned into *SalI* and *KpnI* sites | (6) |
| pGES485 | Integrative vector, Sp<sup>r</sup> | G.E. Schjuman (unpublished) |
| pHPKS | *B. subtilis* low copy number replicative vector, MCL<sup>r</sup> | (38) |
| pJET 1.2/blunt | *E. coli* cloning vector, Amp<sup>r</sup> | Thermo Scientific |
| pJET-*lipL* | *lipL* PCR amplified with oligonucleotides lipL UP and lipL DW cloned in pJET 1.2/blunt | This study |
| pJET-*lipLC150A* | *lipL* C150A PCR amplified with oligonucleotides *lipL_Kpn_FOR* and *lipL_Hind_REV* cloned in pJET 1.2/blunt | This study |
| pJET-*lplJ* | *lplJ* PCR amplified with oligonucleotides *lplJ* UP and *lplJ* DW cloned in pJET 1.2/blunt | This study |
| pJM134 | Integrative vector, Sp<sup>r</sup> | M. Perego (unpublished) |
| pKNT25 | Derived from plasmid pSU40, it allows to create in-frame fusions to the N-terminal end of the T25 fragment of *B. pertussis* adenylate cyclase, Km<sup>r</sup> | (16) |
| pKNT25-*lipL* | pKNT25 containing *lipL* between *XbaI-BamHI* | This study |
| pKNT25-*zip* | Derivative of pKT25 in which a leucine zipper is genetically fused in frame to the T25 fragment | (16) |
| pKT25 | Derived from plasmid pSU40, it allows to create in-frame fusions to the C-terminal end of T25 fragment, Km<sup>r</sup> | (16) |
| pKT25-*lipL* | pKT25 containing *lipL* between *XbaI-BamHI* | This study |
| pNM47 | pJM114 containing *lipL* interrupted with a kanamycin cassette | (6) |
| pNM82 | pBluescript containing a 410 bp fragment of *lipL* gene between *HindIII-ClaI* | This study |
| pNM85 | pSac-Cm containing *lacI Pspac-lplA* cloned into the *sacA* locus | This study |
| pNR002 | pNM82 containing a 846 bp fragment of *lipL* C150A gene in *ClaI* | This study |
| pNR005 | pSG1154 containing *lipL* C150A gene between *KpnI-HindIII* | This study |
| pQC079 | Vector which expresses LipL C150A with a 6-His tag in N-terminus | (8) |
| pSac-Cm | Integrative vector for *SacA* locus, Cm<sup>r</sup> | (36) |
| pSG1154 | Vector used to fuse GFP onto de C-terminus of | (35) |
|  | any protein under the control of a xylose inducible promoter. The fusion is integrated into *B. subtilis amyE* locus |  |
| pUT18 | Derivative of the plasmid pUC19. It allows to create in-frame fusions to the N-terminal end of the T18 fragment of *B. pertussis* adenylate cyclase, Amp<sup>r</sup> | (16) |
| pUT18C | Derivative of the plasmid pUC19. It allows to create in-frame fusions to the C-terminal end of T18 fragment, Amp<sup>r</sup> | (16) |
| pUT18c-lplJ | pUT18C containing *lplJ* gene between *XbaI*-*SacI* | This study |
| pUT18C-zip | Derivative of pUT18C in which a leucine zipper is genetically fused in frame to the T18 fragment, Amp<sup>r</sup> | (16) |
| pUT18-lplJ | pUT18 containing *lplJ* gene between *XbaI*-*SacI* | This study |

<sup>a</sup> Amp, Cm, Km, MCL and Sp denote ampicillin, chloramphenicol, kanamycin, macrolides and spectinomycin resistance cassettes, respectively.

A strain with a deletion of the *lipL* gene was constructed by gene replacement with a kanamycin resistance determinant, through a double crossover event. For this purpose plasmid pNM47 was generated as described previously (6), linearized with *Sca*I and used to transform strain JH642, yielding strain NM28.

For Δ*lipL* complementation and expression analyses, a plasmid expressing LipL C150A was constructed as follows: a 952 bp fragment containing *lipL* with its ribosome binding site (RBS) was PCR-amplified with oligonucleotides LFwSal and LRevBK (6). This fragment was digested with *Hind*III and *Cla*I to obtain a fragment containing the first 410 bp of the *lipL* gene, which was cloned into pBluescriptKS (Stratagene), yielding plasmid pNM82. Plasmid pQC079 (8) was digested with *Cla*I to obtain a fragment of *lipL* gene containing a point mutation that replaces the cysteine 150 for alanine. This fragment was cloned into pNM82 to obtain plasmid pNR002. To construct a translational fusion of the *lipL* C150A gene contained in pNR002 to the green fluorescent protein (GFP), an 860 bp fragment containing *lipL* C150A allele was PCR amplified using oligonucleotides lipL_Kpn_FOR and lipL_Hind_REV. This fragment, cloned in pJET 1.2/blunt (pJET-*lipLC150A*) was digested with *Hind*III and *Kpn*I and inserted into pSG1154 (35), rendering plasmid pNR005. This plasmid was used to transform strain NM28. The double crossover event into the *amy* locus was assessed by the ability to metabolize starch. The resulting strain was named NR008 (*lipL∷*Km^r^ *amyE∷lipL* C150A). A similar strategy, using pGES40-lipL (6) as template, was performed to construct a wild-type LipL-GFP fusion, rendering strain AL107.

To study complementation of Δ*lipL ΔlplJ* strain with *E. coli* LplA, strain NR001 (*lipL∷*Km^r^, *lplJ∷*Sp^r^ *sacA∷*P*spac-lplA*) was constructed. Briefly, wild-type copy of the *E. coli lplA* gene (1055 bp fragment) was PCR-amplified from genomic DNA of strain W3110 with oligonucleotides lplAHFw and lplABRv and the product inserted between the *Hind*III and *BamH*I sites of pGES485 (G. Schujman, unpublished). This plasmid was digested with *Eco*RI and *Bam*HI to obtain a fragment containing *lacI Pspac-lplA*, which was cloned into the *sacA* locus of pSac-Cm (36) previously digested with the same enzymes, yielding plasmid pNM85. Plasmid pNM85 was linearized with *Sca*I and used to transformed strain NM60 (6), yielding strain NM107. This strain was transformed with plasmid pNM47 (6) linearized with *Sca*I, yielding strain NR001. Transformants were screened for *sacA* phenotype, as previously described (36).

For two-hybrid analyses, four plasmids were constructed: pKT25-*lipL,* pUT18*-lplJ*, pKNT25-*lipL* and pUT18C-*lplJ*. These plasmids contain the *lipL* gene fused in frame to the T25 fragment of *Bordetella pertussis* adenylate cyclase and the *lplJ* gene fused in frame to its T18 fragment. Plasmids pKT25-*lipL* and pKNT25-*lipL* were constructed as follows: an 856 bp fragment containing *lipL* gene was PCR-amplified using oligonucleotides lipL Up and lipL DW and ligated into pJET 1.2/blunt to obtain plasmid pJET-*lipL.* This plasmid was digested with *Xba*I and *BamH*I and the resulting fragment was inserted into plasmids pKT25 and pKNT25 (16). Plasmids pUT18*-lplJ* and pUT18C-*lplJ* were constructed as follows: a 1007 bp fragment containing *lplJ* gene was PCR-amplified using oligonucleotides lplJ Up and lplJ DW. This fragment was ligated to pJET 1.2/blunt to obtain plasmid pJET-*lplJ.* This plasmid was digested with *Xba*I and *Sac*I and the resulting fragment was inserted into the plasmids pUT18 and pUT18C (16).

To construct a conditional mutant in the *yloUV* operon a 245 bp fragment containing the natural RBS and the 5’region of *yloU* gene was amplified using oligonucleotides yloU UP and yloU DW. This fragment was inserted into *Hind*III and *Xba*I sites of vector pDH88 (37) rendering plasmid pFM24 (Machinandiarena, personal communication). Strain AL106 was obtained by transformation of strain Δ*lipM* with plasmid pFM24. Integration by a single crossover event results in a truncated copy of the *yloU* gene under its natural promoter and the full *yloUV* operon under the P*spac* promoter. Correct integration of plasmid pFM24 was confirmed by PCR, using oligonucleotides Pspac-1UP and yloU OUT.

The *gcvH odhB* deletion mutant strain CM56, was obtained by transformation of strain NM20 with plasmid pCM1104. This plasmid was constructed as follows: a 2050 bp fragment from the 5′ upstream to the 3’ downstream region of the *odhB* gene was PCR-amplified with oligonucleotides ODHup and ODHdw and cloned in pJET1.2/blunt, yielding plasmid pCM1103. The spectinomycin-resistance cassette from plasmid pJM134 (M. Perego, unpublished) was inserted between the *Hinc*II and *Kpn*I sites of the previously generated plasmid to render plasmid pCM1104.

To obtain a *odhB* deletion mutant strain, chromosomal DNA of strain CM56 was used to transform JH642. Upon selection for spectinomycin-resistance, the colonies that remain sensitive to kanamycin were selected, and the presence of wild type *gcvH* gene was confirmed by PCR (6). This strain was named CM57.

A wild type copy of *odhB* gene was PCR amplified with oligonucleotides odh_Xho_FOR and odh_Cla_REV, and inserted into *XhoI* and *ClaI* sites of plasmid pSG1154 (14), rendering pAL35. The mutant copy of ODHB in which Glu^39^ is replaced by Gln was obtained as follows: oligonucleotides ODHup and ODHExQ_RV were used to amplify the 5’fragment of *odhB* gene while oligonucleotides odhExQ_FOR and odh_Cla_REV, to amplify the 3’end. Both fragments were used as template for an overlap extension PCR in which after 10 cycles of extension, oligonucleotides odh_Xho_FOR and odh_Cla_REV were added. The product obtained was inserted into *Xho*I and *Cla*I sites of vector pSG1154 (35) resulting in plasmid pAL34. Plasmids pAL34 and pAL35 were then digested with *Sal*I and *Cla*I and ligated in vector pHPKS (38) rendering plasmids pAL36 and pAL41, respectively. Both plasmids were used to transform strain CM56.

### Immunoblotting analyses

*B. subtilis* wild type and mutant strains were grown overnight in SMM supplemented with sodium acetate and BCFAP at 37°C. Cells were used to inoculate fresh media of the same composition with or without LA, and cultured at 37°C. After 22 hours of growth, 1 ml aliquot of each sample was centrifuged and the pellets were washed with buffer (20 mM Tris-HCl [pH 8.0], 150 mM NaCl). They were resuspended in 180 μl of lysis buffer (50 mM Tris-HCl [pH 8.0], PMSF 10 mM) per OD_600_ unit. Then, cells were disrupted by incubation with lysozyme (100 μg/ml) for 15 min at 37°C followed by 5 min of boiling in the presence of loading buffer. Each sample was fractionated by sodium dodecyl sulfate-gel electrophoresis in a 12% acrylamide gel. Proteins were transferred to a nitrocellulose membrane and detected using anti-lipoate rabbit antibody and anti-rabbit immunoglobulin G conjugated to peroxidase (Bio-Rad). The bands were visualized using the ECL Plus Western Blotting Detection System (GE).

### Adenylate cyclase two-hybrid assay

The method used for the adenylate cyclase two-hybrid assay was essentially that of Euromedex (16). BTH101 host cells were co-transformed with the following combinations of plasmids: pKT25-*lipL*/pUT18-*lplJ*, pKNT25-*lipL*/pUT18-*lplJ*, pKT25-*lipL*/pUT18C*-lplJ* and pKNT25-*lipL*/pUT18C-*lplJ*. Transformed colonies were grown on LB plates containing 5-Bromo-4-Chloro-3-indolyl-β-D-galactopyranoside (X-gal; 40 μg/ml) and isopropyl β-D-1-thiogalactopyranoside (0.5 mM IPTG) at 30°C for 24 h.

### Fluorescence microscopy

*B. subtilis* NR008 strain was grown overnight in SMM supplemented with sodium acetate and BCFAP at 37°C. Cells were used to inoculate fresh media of the same composition with or without xylose, and cultured at 37°C until they reached exponential growth phase. An aliquot of these cultures was used for microscopy. Microphotographs were taken with a Nikon Eclipse 800 microscope and an Andorclara camera. Exposure time was 30 ms for bright-field microscopy and 5 s for fluorescence microscopy. Images were processed and analyzed with Nis Elements and ImageJ.

### Bioinformatics

Protein sequences were analyzed with the program BLASTP (39). Sequence alignments were performed using T-Coffee (40) and drawn using Boxshade (http://sourceforge.net/projects/boxshade/). The computer program I-Tasser (41) was used to construct a model of the ODH, PDH, BKDH and GcvH lipoyl domains. The models were aligned and visualized in PyMOL (42).

## ACKNOWLEDGEMENTS

We gratefully acknowledge Marta Perego for the gift of plasmid pJM134, and John E. Cronan, Jr., for the gift of pQC079. We acknowledge Diego de Mendoza for critical reading of the manuscript, and Marina Avecilla for technical support. N. Rasetto was a fellow of Consejo Interuniversitario Nacional (CIN), A. Lavatelli is a Fellow of Consejo Nacional de Investigaciones Científicas y Técnicas (CONICET), N. Martin was a fellow of CONICET and M.C. Mansilla is a Career Investigator of the same institution. This work was supported by grants from Agencia Nacional de Promoción Científica y Tecnológica (PICT 2012-1341) and CONICET (P-UE 2016-IBR).

